# MarR-Dependent Transcriptional Regulation of *mmpSL5* induces Ethionamide Resistance in *Mycobacterium abscessus*

**DOI:** 10.1101/2022.10.03.510743

**Authors:** Ronald Rodriguez, Nick Campbell-Kruger, Jesus Gonzalez Camba, John Berude, Rachel Fetterman, Sarah Stanley

## Abstract

*Mycobacterium abscessus* (*Mabs*) is an emerging non-tuberculosis mycobacterial (NTM) pathogen responsible for a wide variety of respiratory and cutaneous infections that are difficult to treat with standard antibacterial therapy. *Mabs* has a high degree of both innate and acquired antibiotic resistance to most clinically relevant drugs, including standard anti-mycobacterial agents. Ethionamide (ETH), an inhibitor of mycolic acid biosynthesis is currently utilized as a second-line agent for treating multidrug resistant tuberculosis (MDR-TB) infections. Here, we show that ETH has activity against clinical strains of *Mabs in vitro* at concentrations that are therapeutically achievable. Using transposon mutagenesis and whole genome sequencing of spontaneous drug-resistant mutants, we identified *marR* (MAB_2648c) as a genetic determinant of ETH sensitivity in *Mabs*. The gene *marR* encodes a transcriptional regulator of the TetR family of regulators. We show that MarR represses expression of MAB_2649 (*mmpS5*) and MAB_2650 (*mmpL5*). Further, we show that de-repression of these genes in *marR* mutants confers resistance to ETH, but not other antibiotics. To identify determinants of resistance that may be shared across antibiotics, we also performed Tn-Seq during treatment with amikacin and clarithromycin, drugs currently used clinically to treat *Mabs*. We found very little overlap in genes that modulate the sensitivity of *Mabs* to all three antibiotics, suggesting a high degree of specificity for resistance mechanisms in this emerging pathogen.

**Importance:** Antibiotic resistant infections caused by *Mycobacterium abscessus* (*Mabs*) have been increasing in prevalence and treatment is often unsuccessful. Success rates range from 30-50%, primarily due to the high intrinsic resistance of *Mabs* to most clinically useful antibiotics. New therapeutic strategies, including repurposing of existing antibiotics, are urgently needed to improve treatment success rates. Here, we show that the anti-TB antibiotic ethionamide (ETH) has repurposing potential against *Mabs*, displaying bacteriostatic activity and delaying emergence of drug resistance when combined with clinically relevant antibiotics currently used against *Mabs in vitro*. We identified genes that modulated susceptibility of *Mabs* to ETH. *marR* encodes a transcriptional regulator that when deleted, confers ETH resistance. Our collective findings can be used to further explore the function of other genes that contribute to ETH susceptibility and help design the next generation of antibacterial regimens against *Mabs* that may potentially include ETH.

## Introduction

*Mycobacterium abscessus* (*Mabs*) is an emerging, non-tuberculosis mycobacterial (NTM) pathogen that has become prevalent in individuals suffering from various underlying lung disorders, including bronchiectasis, chronic obstructive pulmonary disorder (COPD), and cystic fibrosis (1, 2). In the United States, *Mabs* infections represent up to 13% of all NTM pulmonary infections, second only to infections with the *Mycobacterium avium* complex (MAC) (3, 4). The epidemiology of *Mabs* infections has expanded globally in the past decade, representing up to 35% of clinical NTM cases in some regions (5, 6). These infections are difficult to treat with standard antibacterial therapy, with cure rates ranging from 30-50%, making *Mabs* one of the most antibiotic resistant pathogens in clinical settings (7). A better understanding of the biological mechanisms that allow *Mabs* to resist antibiotic action, in combination with drug development and repurposing efforts, is necessary for improving the therapeutic success of *Mabs* infections (8, 9).

Current treatment of *Mabs* infections in healthcare settings requires a minimum of 12 months of intensive antibacterial therapy, which often fails (10). In pulmonary cases, lung resection or transplantation is often necessary (11). Antibiotic treatment options include intravenous or inhaled amikacin (aminoglycoside) in combination with clarithromycin or azithromycin (macrolides), imipenem (β-lactam), cefoxitin (β-lactam), tigecycline (tetracycline), moxifloxacin (fluoroquinolone), and linezolid (oxazolidinone) (10, 12). Resistance to currently administered antibiotics is common and a significant contributor to treatment failure (13). A unique feature of the *Mabs* resistance profile is that the bacteria are intrinsically resistant to most clinically relevant antibiotics (13). However, only a handful of intrinsic resistance mechanisms have been identified to date. Antibiotic-modifying monooxygenases and beta-lactamases that contribute to resistance to tetracycline and various beta-lactam antibiotics have been identified (14, 15). In addition, the transcription factor Whib7 was found to contribute to the expression of genes that confer resistance to numerous ribosome targeting antibiotics (16). While past studies have been able to identify a handful of genes that are important for intrinsic resistance, we still lack a completing understanding of the remarkable breadth of resistance observed in *Mabs*. The *Mabs* genome contains numerous putative phosphotransferases, acetyltransferases, and transporters that could modify and efflux antibiotics (17, 18), but experimental evidence for how these mechanisms relate to specific antibiotics is lacking. It is also unclear whether there are a handful of universal mechanisms that regulate resistance to antibiotics broadly, or whether there are numerous drug-specific resistance mechanisms.

Mycolic acids are known to play critical roles in the pathogenesis, virulence, and impermeability of mycobacteria, and targeting the mycolic acid biosynthetic pathway has yielded several successful anti-mycobacterial agents (19–21). The drug isoniazid (INH) targets mycolic acid biosynthesis by inhibiting the NADH-dependent enoyl ACP reductase InhA, a component of FAS-II (19, 22). INH is the cornerstone of anti-tuberculosis therapy, but has little efficacy against *Mabs* (23–25). The reasons for the lack of efficacy of INH against *Mabs* are unclear (25). Thiacetazone (THZ), an inhibitor of the HadABC dehydratase complex associated with FAS-II, is only modestly active against *Mabs in vitro* but chemical derivatives of THZ have shown improved efficacy (26, 27). These findings suggest that mycolic acid inhibition may represent an untapped research area for developing new drug regimens against *Mabs* infections.

Ethionamide (ETH), a structural analog of INH, is a second-line mycolic acid inhibitor currently used to treat drug-resistant infections associated with *Mycobacterium tuberculosis* (*Mtb*) (28). Similar to INH, ETH inhibits mycolic acid biosynthesis in *Mtb* by inhibiting InhA (23). ETH requires activation by EthA, a flavin-containing monooxygenase, that covalently links NAD to ETH (ETH-NAD) (29, 30). ETH-NAD acts as a competitive inhibitor by competing with NADH for binding in the active site of InhA (30). ETH resistance is common among *Mtb* clinical isolates (31). Mutations in *ethA* reduce ETH bioactivation, conferring resistance (23). Here we show that clinical isolates of *Mabs* are susceptible to ETH in axenic culture at concentrations that are therapeutically achievable *in vivo*. Further, we show that combining ETH with other clinically used antibiotics results in suppression of resistance emergence.

Tn-Seq is a powerful tool that facilitates the identification of genes important for bacterial growth under any condition in a high-throughput manner (32). Tn-Seq has been previously used to identify genes important for growth during antibiotic exposure in *Pseudomonas aeruginosa, Staphylococcus aureus, and Mycobacterium avium* (33–35). Tn-Seq libraries have been previously generated in *Mabs*, and used to explore essential genes for growth in axenic culture, as well as genes required for survival in a lung epithelial model (36–38). Here, we use Tn-Seq to identify transposon mutants that display both growth defects and growth advantages when exposed to the ETH. We identify a *Mabs marR* homolog as an important genetic determinant of sensitivity to ETH. Point mutations and deletion of *marR* confer ETH resistance. This increase in resistance is due to upregulation of *mmpL5 and mmpS5*, which encodes for two putative transport proteins that are part of two larger families of proteins (MmpS and MmpL proteins) known to confer resistance to other antibiotics in *Mabs* (27, 39). Deletion of these genes re-sensitizes ETH-resistant bacteria. Interestingly, upregulation of *mmpL5* and *mmpS5* does not appear to confer broad antimicrobial resistance. Indeed, comparison of Tn-Seq results across the antibiotics ETH, amikacin, and clarithromycin reveals a surprising lack of overlap, suggesting a high degree of specificity in mechanisms that modulate antibiotic susceptibility/resistance in this important pathogen. Taken together, this work suggests that ETH may be of interest for treating drug resistant *Mabs* infections in combination with other effective antibiotics.

### Results

### Transposon mutagenesis demonstrates essentiality of the mycolic acid biosynthetic pathway in Mabs

To identify essential genes in *Mabs* on a genome-wide scale we constructed a transposon mutant library in WT *Mabs* ATCC 19977 by packaging the *Himar1* transposon in phage ΦMycoMarT7 followed by bacterial transduction (40). We chose this strain since it has a fully sequenced genome and is genetically tractable (41, 42). To determine the location and abundance of transposon insertions in the resulting library, we collected genomic DNA and sequenced the transposon junctions (43, 44). We obtained approximately 80,000 unique transposon insertions in non-essential genes. We were not able to obtain insertions in 376 genes, suggesting that these genes are essential. These data are largely consistent with previous studies that used transposon mutagenesis to define the essential genome in *Mabs* (36, 37).

Genes associated with translation, ribosomal structure, and biogenesis were the most represented cluster of orthologous group (COG) category among those without insertions (Fig. 1A). Interestingly, only 51% of the predicted essential genome of *Mabs* is shared with *Mtb* (Fig. 1B). Of the 376 predicted essential genes, 57 genes (approximately 15.2%) do not have obvious homologs in *Mtb*. For 18 genes known to play a role in mycolic acid biosynthesis in *Mycobacterium tuberculosis*, most putative homologs in *Mabs* were also found to be essential in our analysis (Table S1) (45, 46). Some of these genes include *inhA* (MAB_2722c), *fadD32* (MAB_0179), *pks13* (MAB_0180), and *mmpL3* (MAB 4508) (Table S1) (45). The products of the genes have been used to develop various chemical inhibitors of mycolic acid biosynthesis in *Mtb* (19). The potential of this shared essentiality in *Mabs* suggests that the mycolic acid pathway may also be vulnerable to drug targeting.

**Figure 1.**
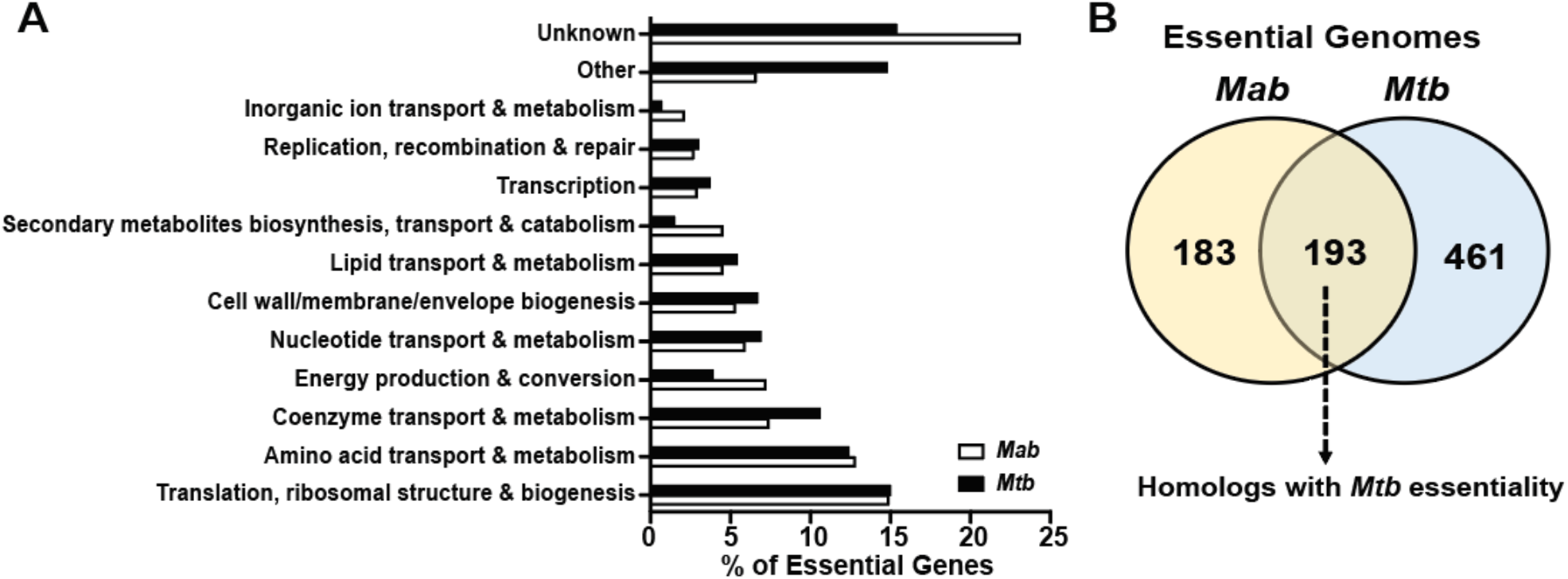
Tn-Seq analyses of the essential genome in *Mabs*. (A) Essential genes in *Mabs* were grouped in cluster of orthologous groups (COG) and then compared to the essential genome in *Mtb*. ‘Other’ represents COG categories containing less than 2% of the essential genome. ‘Unknown’ represents genes without any COG annotations. (B) Venn diagram displaying essential gene homologs in *Mabs* and *Mtb*.

### Mycolic acid inhibitors are largely ineffective against Mabs, although ETH activity displays promising repurposing potential

To test antimicrobial activity of mycolic acid inhibitors against *Mabs*, we determined the minimum inhibitor concentration (MIC) of representative mycolic acid inhibitors: isoniazid (INH), thiacetazone (THZ), and ETH (23, 26, 27), against WT *Mabs*. We chose INH and ETH because they possess clinical relevance in the context of *Mtb* and their mechanism of action has been extensively studied (23, 31, 47). The MIC_99_ for INH (1.25 mg/ml) and THZ (> 500 μg/ml), are much higher than those commonly observed with drug sensitive *Mtb* and are above standard clinical breakpoints (Fig. 2 A-B, Table S3) (48, 49). The high-level of intrinsic resistance displayed by *Mabs* to these agents is a major reason why they are not used in clinical settings where *Mabs* infections are observed. However, unlike other mycolic acid targeting drugs, we found that the ETH MIC_99_ (8 μg/ml) is below a concentration that can be achieved therapeutically (20 μg/ml) (Fig. 2C) (50). Indeed, we found that ETH at this concentration has modest bactericidal activity against *Mabs* (Fig. 2D). Importantly, the ETH MIC of 9 clinical isolates tested were similar to our WT ATCC 19977 strain, suggesting that relative sensitivity to ETH may not be a unique feature of a single strain (Fig. 2E). This observation suggests that sensitivity to ETH at therapeutically relevant levels is a common feature of *Mabs* clinical isolates.

**Figure 2.**
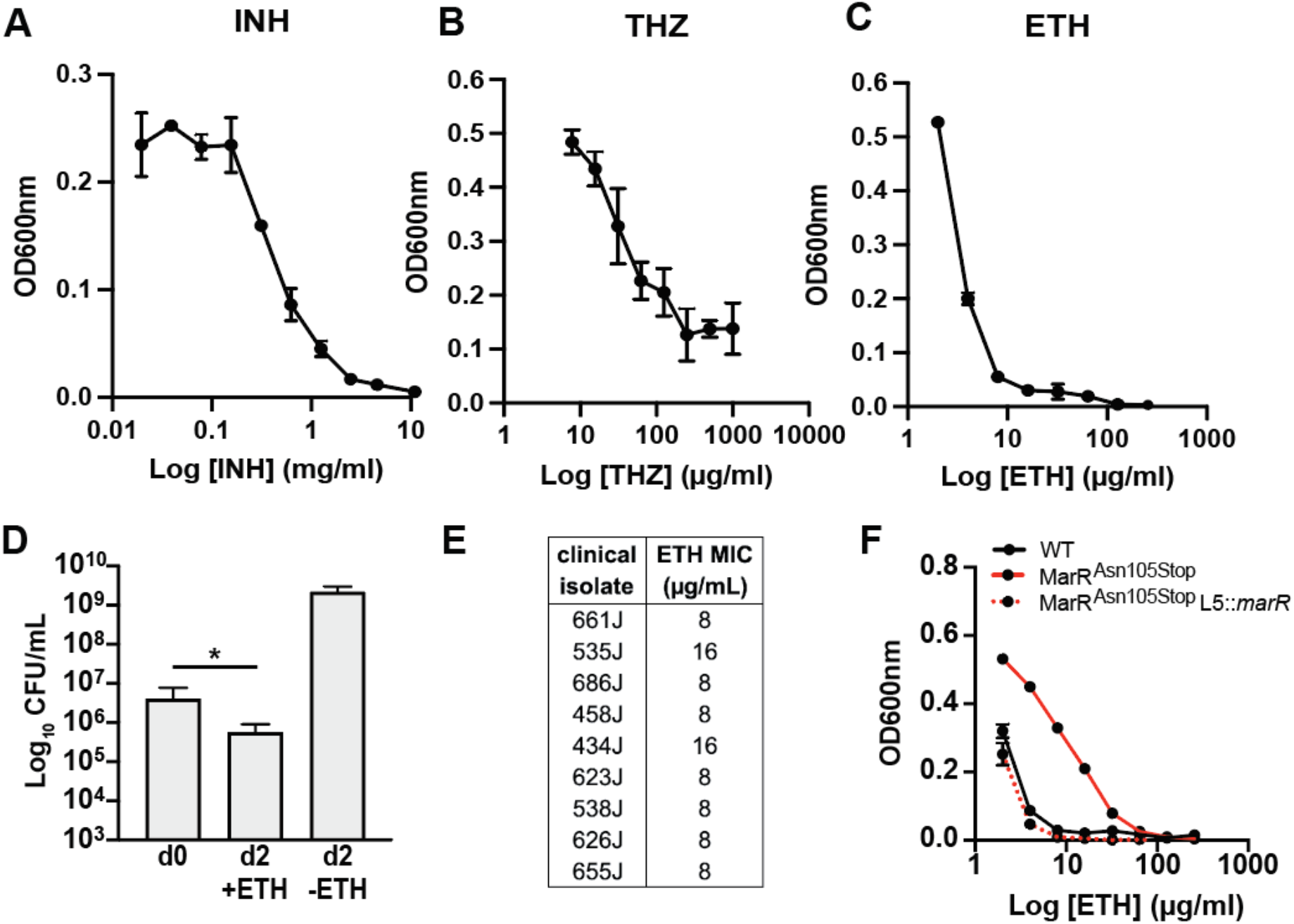
ETH displays mild bactericidal activity against *Mabs*. Dose-response curves for (A) INH, (B) THZ, and (C) ETH against WT *Mabs*. (D) WT *Mabs* was treated with and without ETH (20 μg/ml) for 2 days and viable bacteria were enumerated by CFU (E) ETH MIC values for clinical isolates of *Mabs*. (F) ETH dose-response curves for WT, MarR^Asn105Stop^, and MarR^Asn105Stop^ complement (MarR^Asn105Stop^ L5::*marR*) strains. All experiments are representative of at least three biological replicates except for (E), which is representative of two biological replicates. Error bars represent standard deviation, *p* ≤ 0.05 (*).

### Whole genome sequencing of ETH-resistant bacteria identifies mutations in marR (MAB_2648c)

In *Mtb*, ETH resistance in drug-resistant clinical strains is often conferred by mutations in *ethA* or *inhA* (23). To identify mutations that confer resistance to ETH in *Mabs*, we generated spontaneous resistant mutants by plating WT bacteria on solid media containing twice the MIC of ETH for *Mabs* (Fig. S1A). Three resistant isolates (R128, R150, and R200) were selected for further analysis. First, to validate that the isolated mutants were indeed ETH resistant, we exposed WT and mutant bacteria to a two-fold dilution series of ETH in broth. We found that the mutants were approximately 8-fold more resistant (8 μg/ml in WT and 64 μg/ml in all spontaneous clones) to ETH than WT (Fig. S1B-D). To identify the genetic basis for resistance, we performed whole genome sequencing of R128, R150, and R200. Only one mutation was common across all three mutants, consisting of a CG dinucleotide deletion in MAB_2648c at nucleotide positions 354 and 355. This deletion alters the reading frame of this gene, introducing a premature stop codon at amino acid position 105. MAB_2648c encodes for a putative *marR* (**m**ultidrug **a**ssociated **r**esistance **r**egulator) transcriptional regulator that represents a subtype of TetR transcriptional repressors, many of which have been associated with resistance to functionally diverse antibiotics in different bacteria (51). Introducing an integrative copy of WT *marR* in the spontaneous mutants restored ETH sensitivity (Fig. 2F). Many clinical isolates of *Mtb* that display acquired resistance to ETH are cross-resistant to INH due to mutations in the promoter of *inhA* (52). Our isolated *marR* mutants do not have increased resistance to INH relative to the parental wild-type strain (Fig. S1B-D).

### Transposon insertions in marR confer a growth advantage in the presence of ETH

To more systematically identify additional genes that modify the efficacy of ETH against *Mabs*, we screened mutants in our Tn library for changes in growth in the presence of a subinhibitory ETH concentration (Fig. 3A, S2A). The Tn library was cultured in the presence of ETH for 24 hours, at which time surviving bacteria were plated on agar plates. DNA was prepared from resulting colonies and transposon gene junctions were amplified and sequenced as previously described (43, 44); changes in transposon abundance after exposure to ETH were analyzed using TRANSIT (Table S2) (44). We identified 208 genes as potential modulators of susceptibility to ETH using a two-fold change cutoff and a *p-*value of ≤0.05 (Fig. 3A). Among genes with annotations, those associated with transcription were the most abundant, representing approximately 9% of the identified hits. (Fig. S2B). Interestingly, mutations in two of the three *ethA* homologs in *Mabs* seemed to confer resistance in the screen (Fig. 3A), suggesting that multiple monooxygenases may contribute to ETH activation in *Mabs*.

**Figure 3.**
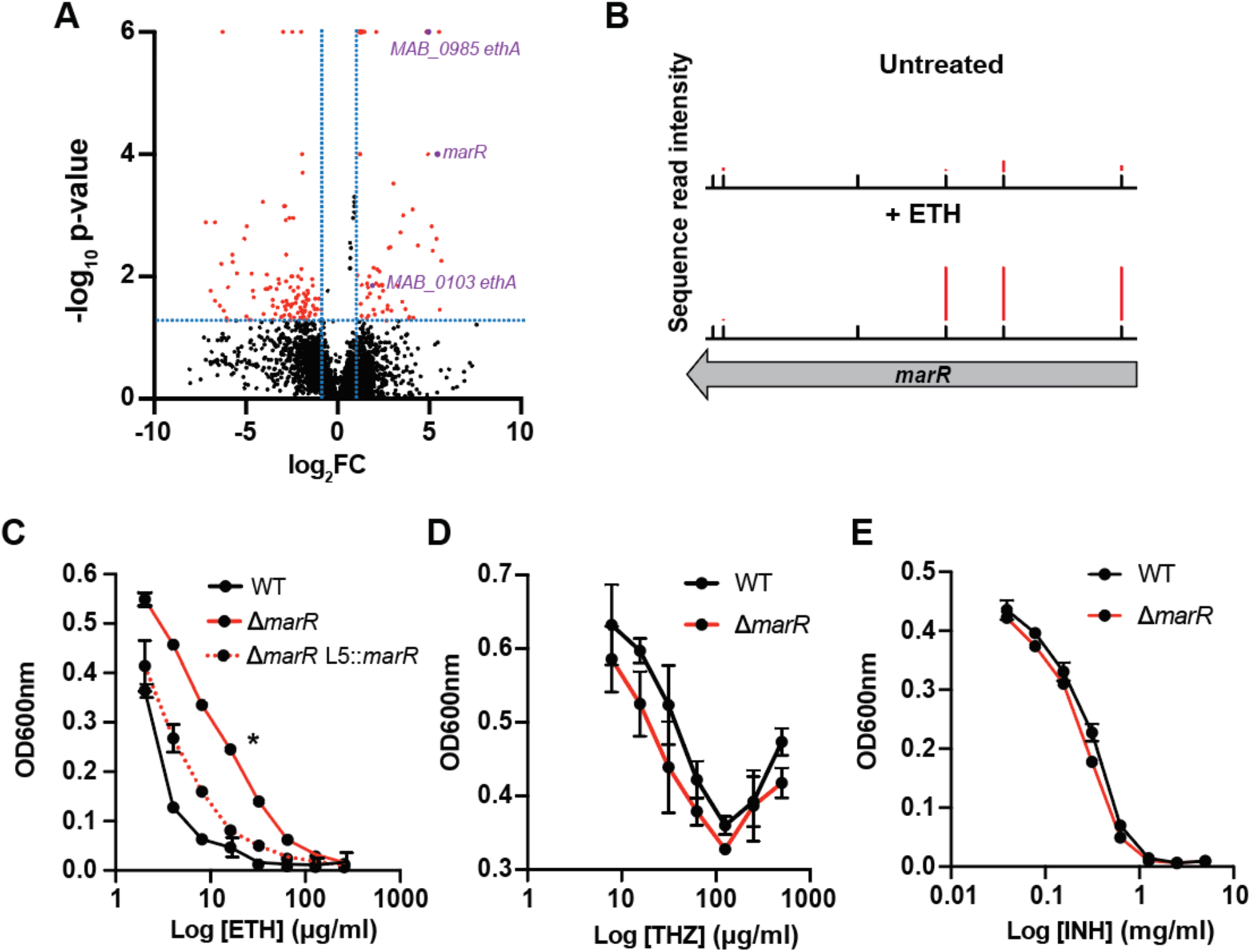
*marR* is a determinant of ETH susceptibility. (A) Volcano plot displaying all transposon insertion mutants from a Tn-Seq screen in the presence of ETH. Red dots represent mutants with log fold change >2 and p<0.05 (B) Transposon sequence read densities for *marR* in the presence and absence of ETH. Black tick marks represent all possible insertion sites. (C-E) Dose-response curves for WT, Δ*marR*, and Δ*marR* complement (Δ*marR* L5::*marR*) strains in the presence of ETH (C), THZ (D), and INH (E). All experiments are representative of at least three biological replicates. Error bars represent standard deviation. *p* ≤ 0.05 (*) for Δ*marR* vs complemented strain.

In agreement with our data from the generation of spontaneous resistant mutants, we found that bacteria carrying insertions in *marR* were approximately five-fold more abundant in the presence of ETH (Fig. 3A-B). To validate these findings, we constructed a deletion mutant of *marR* (Δ*marR*) and examined growth in the presence and absence of ETH. We found that Δ*marR* was approximately 8-fold more resistant to ETH compared to WT bacteria (8 μg/ml for WT and 64 μg/ml for Δ*marR*) (Fig. 3C). Thus, using two independent methods, we show that loss of MarR function leads to ETH resistance. Deletion of *marR* did not confer resistance to any other antibiotic tested, including INH and THZ (Fig. 3D-E) (Table S3). This result is surprising, as in other bacteria, including *E. coli* (single MarR regulator) and *P. aeruginosa* (13 MarR regulators), loss of MarR activity confers resistance to a diversity of antibiotics (53–55).

### MarR negatively regulates expression of mmpS5-mmpL5 and contributes to ETH resistance

TetR regulators often negatively regulate expression of target genes that are transcribed divergently from the regulator, separated by an intergenic region of approximately 200 base pairs (51). The genes *mmpS5* (*MAB_2649*) and *mmpL5* (*MAB_2650*) are in an operon located 369bp upstream of *marR* (Fig. 4A). Mutations in TetR regulators often lead to upregulation of *mmpS* and *mmpL* genes, which contribute to clofazimine, bedaquilline, and THZ resistance in *Mabs* (27, 39). We hypothesized that *mmpS5* and *mmpL5* contribute to ETH resistance, and that in WT bacteria their expression is repressed by MarR. To address this hypothesis, we first measured expression levels of *mmpS5* and *mmpL5* by qRT-PCR in WT and Δ*marR* bacteria. Regardless of the presence of ETH, expression levels of *mmpS5* were upregulated >80-fold in Δ*marR* compared to WT (Fig. 4B). A similar trend was observed for *mmpL5* (Fig. 4C). The addition of ETH to bacterial cultures alone did not affect expression of *mmpS5* (Fig. 1D) or *mmpL5* (Fig. 1E) in WT bacteria, suggesting that MarR activity is not directly regulated by ETH. To determine whether upregulated expression of *mmpS5* and *mmpL5* in Δ*marR* contribute to ETH resistance, we overexpressed both genes in WT bacteria under an anhydrotetracycline (ATc) inducible promoter (56) and examined bacterial growth in the presence of ETH. Overexpression of *mmpS5* and *mmpL5* increased the ETH MIC (> 16 μg/ml) compared to bacteria not induced with ATc (16 μg/ml) (Fig. 4F). These data suggest that upregulation of *mmpSL5* is responsible for the ETH resistance phenotype observed in the absence of *marR*. To further test this finding, we investigated the contribution of *mmpL5* and *mmpS5* to ETH resistance in MarR^Asn105Stop^ bacteria. In agreement with our Δ*marR* qPCR data, we also found that *mmpSL5* is expression is upregulated in MarR^Asn105Stop^ bacteria compared to WT (Fig. S3 C-F). Deletion of *mmpSL5* (Δ*mmpSL5)* in this background sensitized bacteria to ETH at least four-fold compared to MarR^Asn105Stop^ bacteria (4 μg/ml in Δ*mmpSL5* and 16 μg/ml in MarR^Asn105Stop^ bacteria) (Fig. 4G). These results suggest that the loss of MarR activity leads to upregulation of *mmpSL5*, whose activity ultimately provides ETH resistance.

**Figure 4.**
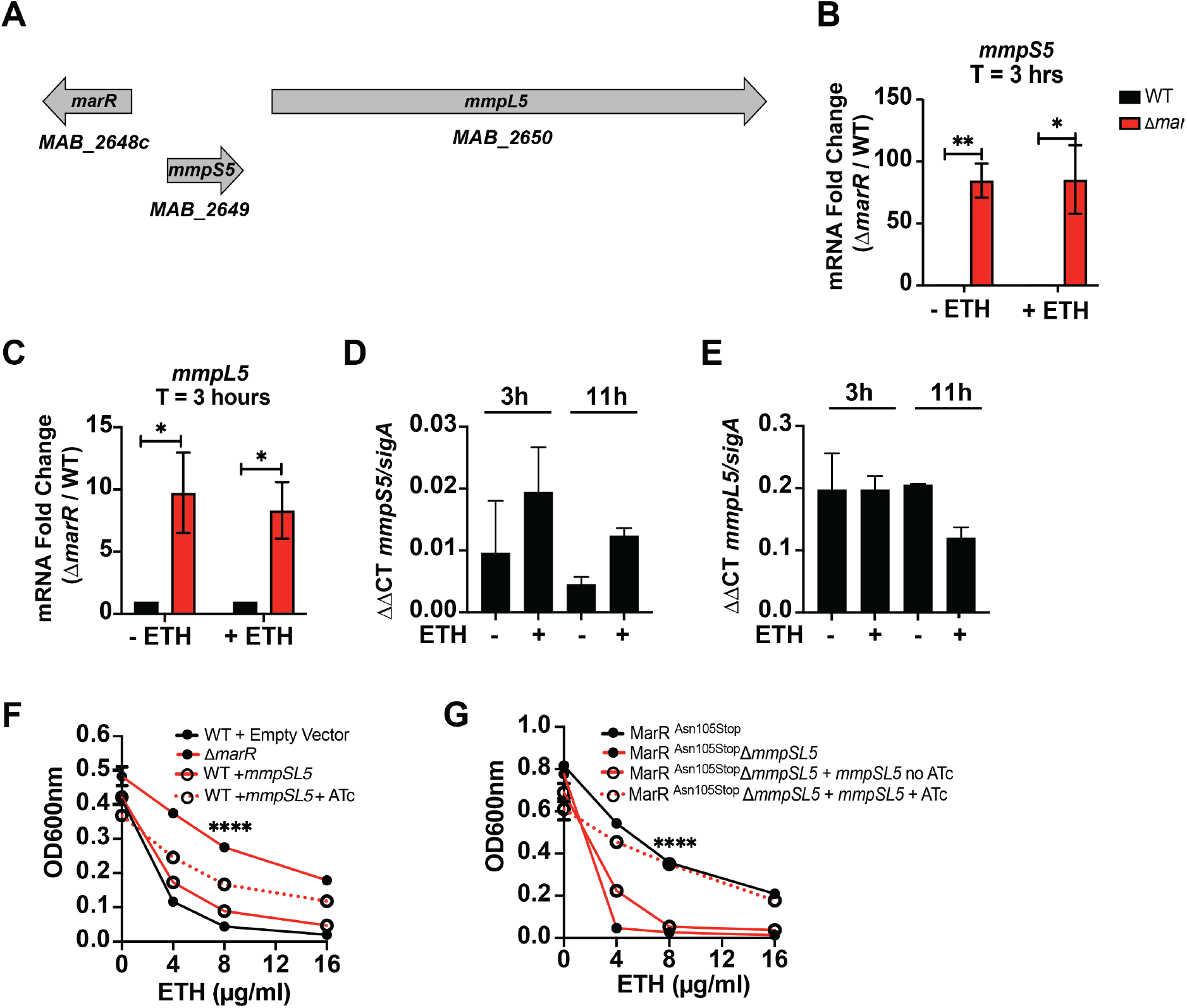
*mmpS5* and *mmpL5* are negatively regulated by MarR and contribute to ETH resistance. (A) Schematic representation of the genetic organization of *marR, mmpS5*, and *mmpL5*. (B-E) Gene expression levels of *mmpS5* and *mmpL5* in WT and Δ*marR* bacteria in the presence and absence of ETH at the indicated time points. (F-G) ETH dose-response curves for the indicated strains. ATc (100 ng/ml) was included where appropriate. All experiments were performed at least in biological duplicates. Error bars represent standard deviation. *p* ≤ 0.05 (*); *p* represent standard deviation. p ≤ 0.01 (**); *p* ≤ 0.0001 (****) comparing *mmpSL5* mutant with and without ATC.

We hypothesized that MmpSL5 may function as a transporter that exports ETH, because many membrane transporters function as efflux pumps to export antibiotics (57). To test this hypothesis, we tested whether the Δ*marR* mutant strain that expresses *mmpSL5* at high levels displays higher rates of ethidium bromide (EtBr) efflux, an assay commonly used to assay mycobacterial efflux in the context of drug resistance (58). Although we observed EtBr accumulating in *Mabs* cells in a dose dependent manner (Figure S4), we did not observe any difference in EtBr levels when comparing WT and the Δ*marR* mutant at a single dose. The inability to detect increased EtBr efflux in the Δ*marR* mutant does not rule out the hypothesis that *mmpSL5* is a transporter of ETH, but it suggests there may be specificity to the transport that precludes the use of this assay.

### ETH treatment with CLR or AMK suppresses the emergence of resistant mutants

Because *Mabs* infections are almost never treated with sole antibiotic regimens (10), we next tested whether ETH synergizes or antagonizes with commonly prescribed antimicrobial agents. We tested ETH in combination with three front line drugs for treating *Mabs* infections that are representative of antibiotics with different mechanisms of action: amikacin (AMK, aminoglycoside), clarithromycin (CLR, macrolide), and moxifloxacin (MFX, fluoroquinolone). Treatment with ETH (2.5X MIC) or MFX (3X MIC) alone resulted in mildly bactericidal activity during the first 2 days of treatment (Fig. 5A). However, after day 2 bacteria grow in the presence of either antibiotic (Fig. 5A), likely due to the emergence of resistant mutants. Although we did not observe either synergy or antagonism at early timepoints of treatment with ETH and MFX after 2 days of treatment, there was extensive bacterial growth after day 2, even in the presence of both antibiotics (Fig. 5A). Similar to MFX, treatment with AMK (12.5X MIC) or CLR alone (0.75X MIC) resulted in the rapid emergence of resistant mutants. However, the combination of ETH with either AMK or CLR prevented the outgrowth of bacteria that occurred after 2 days of treatment with single agents (Fig. 5B-C). These data suggest that ETH may have the potential to delay resistance emergence to clinically relevant antibiotics used against *Mabs*.

**Figure 5.**
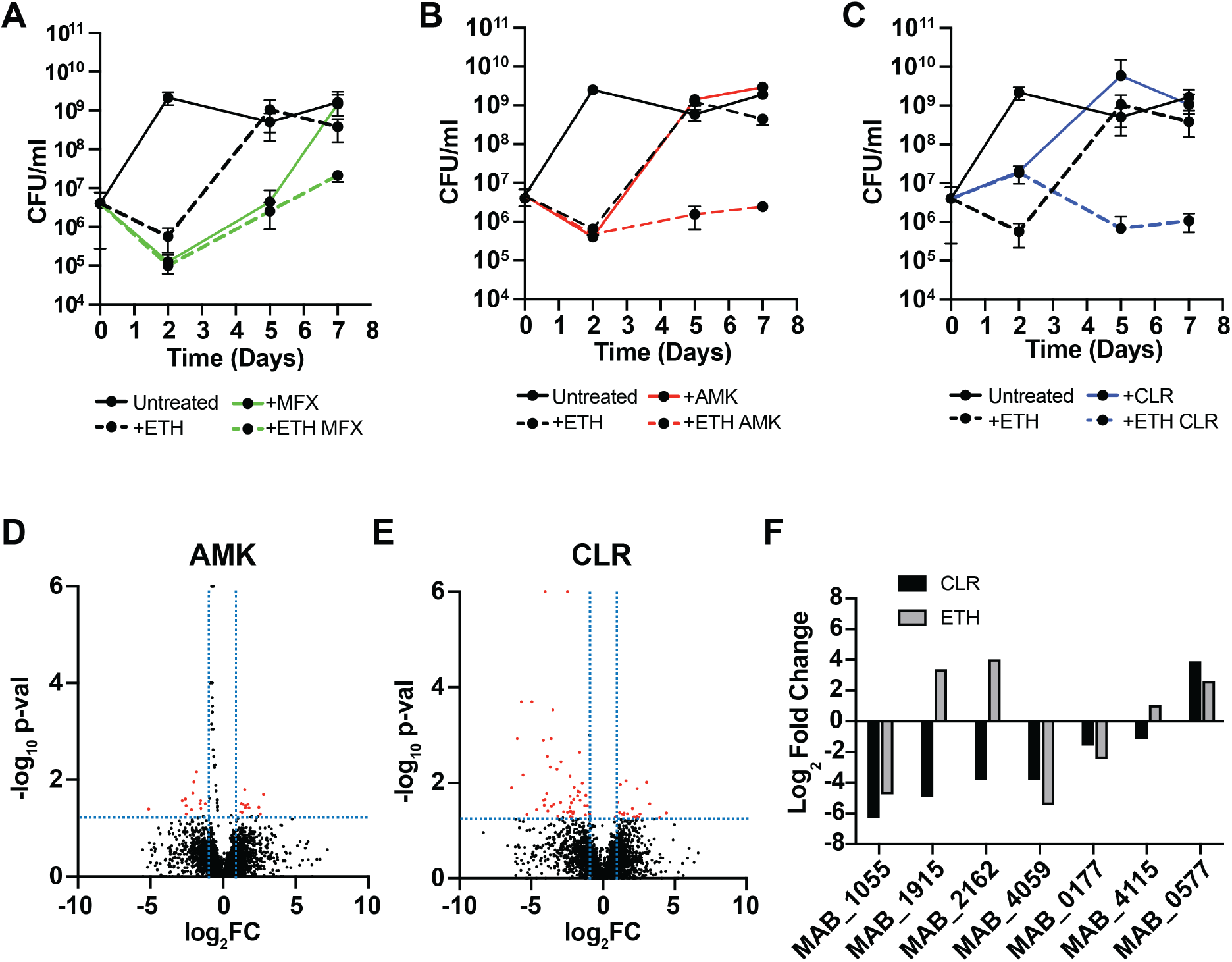
Prolonged ETH treatment suppresses emergence of drug resistance to some clinically relevant antibiotics. (A-C) WT *Mabs* was treated with and without ETH (20 μg/ml) in the presence of moxifloxacin (MFX, 3 μg/ml), amikacin (AMK, 25 μg/ml), and clarithromycin (CLR, 3 μg/ml) or in combination with ETH. At the indicated time points, bacteria were collected, washed, and then ten-fold serial dilutions were plated on LB agar for CFU/ml enumeration. (D-E) Volcano plots displaying all gene hits from a Tn-Seq screen in the presence of AMK (D) and CLR (E). Red dots represent mutants that display growth advantages while black dots represent mutants that display growth defects in the presence of the indicated drug. (F) Fold changes in the transposon read intensities of select mutants from our Tn-Seq screen with ETH and CLR. All experiments were performed in biological duplicates. Error bars represent standard deviation. no significance (ns); *p* < 0.05 (*)

Δ*marR* did not exhibit cross resistance to other antibiotics tested (Table S3), suggesting a unique function in mediating ETH resistance. This was surprising, as MarR homologs often confer resistance to multiple antibiotics in other bacterial species (53–55). The fact that resistant mutants fail to emerge in the combinatorial treatment of ETH with AMK and CLR suggests that there may be unique mechanisms of resistance to each of these antibiotics. To determine whether there are mediators that confer susceptibility or resistance across multiple antibiotics in *Mabs*, we performed TnSeq in the presence of both AMK and CLR as described for ETH (Table S2). Using the same fold-change and significance cutoffs as in our ETH Tn-seq, we found 28 significant genes in AMK and 70 genes in CLR (Fig. 5D-E, S5, S6). No genes were significant in both the ETH and AMK conditions, and only 7 genes were important in both the ETH and CLR conditions. The genes significant under both ETH and CLR are *MAB_1055* (Conserved Hypothetical Peptidase), *MAB_1915* (Probable Fatty Acid CoA Ligase FadD), *MAB_2162* (Putative AAA-family ATPase Mpa), *MAB_4059* (Hypothetical Protein), *MAB_0177* (Antigen 85-A/B/C Precursor), *MAB_4115* (Putative MmpL Membrane Protein), and *MAB_0577* (Putative ABC Transporter Solute Binding Protein). Insertions in three of these genes conferred a growth defect in both conditions, while insertions in only one gene conferred a growth advantage in both conditions (Fig. 5E). Insertions in the remaining three genes had differential effects in the two conditions.

## Discussion

*Mabs* is remarkable among drug-resistant bacterial pathogens in that resistance is intrinsic but can also be acquired to functionally diverse antibiotics (17). Unlike most antibiotics that target mycolic acid biosynthesis, we show here that ETH is mildly bactericidal against *Mabs in vitro* and has the potential to be repurposed with antibiotics currently used against *Mabs* infections. Using a combination of bacterial genetics, whole genome sequencing, and transposon mutagenesis, we identified *marR* as an important determinant of ETH sensitivity in *Mabs*, as loss of *marR* leads to ETH resistance. This loss is associated with upregulation of *mmpSL5*, which contributes to ETH resistance in a *marR*-dependent manner. Our findings expand on the current knowledge regarding the role of *mmpSL* genes in conferring antibiotic resistance in *Mabs*. In line with what we have reported here, the loss of other TetR regulators in *Mabs* leads to upregulation of other *mmpSL* genes that confer resistance to bedaquiline, clofazimine, and thiacetazone (27, 39). It speculated that that many of the *mmpSL* genes in *Mabs* encode for transporters that efflux these antibiotics into the extracellular space (18), although this requires further evaluation. Whether ETH is being transported by MmpSL5 remains unknown. Interestingly, the TetR regulator *Mab_2299c* negatively regulates two, genetically distant *mmpSL* couples (*MAB_2300-2301* and *MAB_1135c-1134c*) that confers cross-resistance to bedaquiline and clofazimine (39). We did not find any differences in bedaquiline and clofazimine susceptibility in the absence of *marR* (Table S3). TetR regulators represents the largest class of transcriptional regulators in *Mabs*, a feature that is characteristic of saprophytic mycobacteria (51). Of the 139 TetR regulators found in the *Mabs* genome, 21 of these are annotated as MarR (59). It is possible that the other MarR regulators confer resistance to antibiotics with diverse modes of action, although this remains to be explored.

We now have several examples of TetR regulators conferring drug resistance in *Mabs*, highlighting the importance of deciphering the roles of those that have yet to be studied in the context of drug resistance (27, 39). Many of these findings have been obtained through the generation of laboratory, drug-resistant strains and have yet to be extended to clinical isolates (27, 39). Sequencing of TetR regulators in drug-resistant clinical isolates could help us identify novel genetic determinants of drug resistance. In our ETH Tn-Seq screen, we identified transposon (Tn) mutants with insertions in 17 TetR regulators (Table S2). Tn mutants with insertions in three other genes besides *marR* (*MAB_2885, MAB_2731*, and *MAB_0979*) displayed a growth advantage in the presence of ETH while mutants with insertions in the other 13 TetRs displayed a growth defect (Table S2). In our Tn-Seq screen with AMK, we identified Tn insertions in 2 TetR regulators: *MAB_2061c* (growth defect) and *MAB_4026c* (growth advantage) (Table S2). Mutants with Tn insertions in two additional TetR regulators (*MAB_1881c* and *MAB_2952c*) displayed growth defects in our Tn-Seq screen with CLR (Table S2). These Tn-Seq hits can be utilized to expand our knowledge on the contribution of TetR regulators as being important genetic determinants for growth on clinically relevant antibiotics.

Many questions regarding the role of MarR in ETH resistance remain. TetR regulators can be dissociated from DNA through direct binding of small ligands (51). For MarR regulators, many of these ligands are aromatic compounds (51, 60). It remains unknown what biological conditions lead to transcriptional de-repression of MarR targets, leading to upregulation of *mmpSL5* and possibly other genes. We speculate that these conditions are associated with ETH resistance, given that ETH exposure alone does not induce expression of either *marR* or *mmpSL5* (Fig. 4B-E, Fig. S3A-B), suggesting that MarR may be associated with functions unrelated to drug resistance (51, 60, 61). The closest protein homologues of Mab_2649 and Mab_2650 in Mtb H37Rv are MmpS5 (Rv0677c, 38.57%) and MmpL5 (Rv0676c, 48.56%), respectively (62). MmpSL transporters are known to transport many lipids, some of which include TMM (MmpL3), PDIM (MmpL7), and sulfolipids (MmpL8) (63). MmpS/L4 and MmpL5 are associated with mycobactin and carboxymycobactin export, which are two of the major mycobacterial siderophores that allow for iron acquisition (63, 64). *Mtb* mutants defective in these transporters are unable to grow in low-iron environments (64). Although there are two MarR regulators in *Mtb* (Rv0042c and Rv0880), these do not share any homology with the MarR regulator identified in this work, suggesting that the mechanism by which MarR contributes to ETH resistance may be distinct from what is currently known in *Mtb* (59, 62).

Identifying overlapping hits in our Tn-Seq screens with CLR and ETH was surprising. While ETH is known to target cell envelope metabolism in *Mtb*, this remains to be determined in *Mabs*. CLR synergizes with the cell wall targeting antibiotic vancomycin against *Mabs in vitro* (65), although the mechanisms responsible for this synergy remain unknown. We found that a combination of CLR and ETH was indifferent against *Mabs* (Table S4). Interestingly, Tn insertions in genes that conferred a growth disadvantage in the presence of CLR and ETH independently (*MAB_1055* and *MAB_0177*) encode for proteins that are predicted to localize to the cell envelope (Fig. 5F). It is possible that a combinatorial action of CLR and ETH can lead to cell surface changes that makes the mycobacterial envelope more permeable to antibiotic entry. Although we did not observe synergy between CLR and ETH against WT *Mabs*, examining this combinatorial action in the absence of *MAB_1055* and *MAB_0177* may display a synergistic effect.

Our collective findings in this work can be used to broaden our knowledge on identifying genetic determinants of ETH and other clinically relevant antibiotics. Identifying a gene that seems to uniquely confer resistance to ETH, but not other unrelated antibiotics, suggest that intrinsic drug resistance in *Mabs* may result from a multitude of genetic mechanisms. The possibility of using ETH in clinical settings should be further explored in the future using mouse models of infections and high-throughput chemical screening experiments. Identifying chemical compounds that synergize with ETH both *in vitro* and *in vivo* along with identifying other mechanisms of ETH resistance in *Mabs* should be the focal point of future work.

## Materials and Methods

### Construction of bacterial strains and growth conditions

*Mabs* strain ATCC 19977 was grown in Middlebrook 7H9 (BD) media supplemented with albumin-dextrose-saline (ADS). For growth on solid media, *Mabs* was cultured on LB agar (BD) or Middlebrook 7H10 (BD) where indicated. Clinical strains were obtained from Chao Qi (Northwestern University). The following antibiotics were supplemented when appropriate for *Mabs*: hygromycin B (Invitrogen)(125 μg/ml for liquid media; 1 mg/ml for solid media), zeocin (InvivoGen)(50 μg/ml), and kanamycin (Sigma)(150 μg/ml for liquid media; 100 μg/ml for solid media). To favor the generation of S morphotype *Mab*s, both liquid and solid media were supplemented with Tween-80 (Sigma, 0.05% v/v). *E. coli* was grown in LB supplemented with the following antibiotics when appropriate: hygromycin B (125 μg/ml), zeocin (50 μg/ml), and kanamycin (50 μg/ml). Deletion of *marR* and *mmpSL5* was achieved using Oligonucleotide-Mediated Recombineering followed by Bxb1 Integrase Targeting (ORBIT) as previously described (66). We adapted the original ORBIT protocol for utilization in *Mabs*. Briefly, for each target gene, an oligonucleotide containing an *attP* site flanked by 60-80 base pairs of sequence homology surrounding each target gene was constructed and co-electroporated (2.5 kV, 25 μF, and 1000 Ω) with the payload plasmid pKM496 into bacteria expressing genes from the plasmid pKM444 (66). All electroporations were conducted with 385 μl of bacteria washed three times with 10% glycerol at an optical density of 600 nanometers (OD_600_) of 0.1-0.3. Approximately 67 ng of target oligonucleotides (Integrated DNA Technologies) were used with and without 100 ng of pKM496 for electroporations using cuvettes containing 0.2 cm gaps (Bio-Rad). Following electroporation, bacteria were rinsed with 2 ml of 7H9 and allowed to recover at 37°C with shaking for 16 hours before plating on LB supplemented with zeocin (50 μg/ml) to select for deletion mutants. PCR amplification was used to check for the correct recombination event, which was confirmed with Sanger sequencing.

### Tn-Seq screening

The transposon (Tn) library for *Mabs* was constructed as previously described for other mycobacterial species (40). For screening, an aliquot of the *Mabs* Tn library was grown in 150 ml of 7H9 broth until log phase (OD_600_ 0.5-1.0). The Tn library was diluted to an OD_600_ of 0.01 (∼ 5 × 10^6^ CFU/ml) in 10 ml of 7H9 with and without ETH (TCI Chemicals, 16 μg/ml), CLR (Ambeed, 1 μg/ml), and AMK (Ambeed, 8 μg/ml). All cultures were prepared in triplicate. Bacteria were grown at 37°C for 24 hours, harvested by centrifugation, and washed twice with fresh 7H9 broth. 2 ml of bacteria containing 1.5 × 10^4^ CFU/ml were plated on LB agar supplemented with kanamycin (100 μg/ml) and Tween-80 (0.05 %) on 245 mm square bioassay dishes (Corning). Plates were incubated at 37°C for 5 days and then colonies were scraped into 35 ml of 7H9 broth. Genomic DNA was extracted from the collected mixture as previously described. Genomic DNA was submitted to the UC Davis DNA Technologies Core and sequences enriched for the *Himar1* transposon were amplified on a HiSeq illumina platform (43). Sequence reads were mapped to the *Mabs* ATCC 19977 genome. Transposon-enriched sequence reads were then analyzed using TRANSIT software (44).

### Determination of Minimum Inhibitory Concentrations (MIC) and synergy testing

*Mabs* strains were grown to an OD600nm of 0.2-1.0 in 7H9 and diluted to an OD600nm of 0.01 with and without a two-fold dilution series of antibiotics to be tested in 96 well, TC-treated plates in a final volume of 100 μl. Plates were incubated at 37°C for four days without shaking in tightly sealed, moist Tupperware containers to prevent evaporation. After 4 days, bacteria were fixed with an equal volume of 5% Formalin (Sigma) and optical densities were recorded at 600 nm using a SpectraMax M3 Microplate Reader (Molecular Devices). The MIC here is defined as the lowest concentration of antibiotic that inhibits 99% of bacterial growth. For synergy experiments, ETH MIC was determined as described above with and without the highest concentration of each antibiotic tested that has no effect on bacterial growth. For colony forming unit (CFU) enumeration, bacteria were washed twice in antibiotic-free growth media and then ten-fold serial dilutions were prepared followed by plating on LB agar. CFU/ml was enumerated after 5 days of incubation at 37°C.

### Selection of spontaneous ETH resistant mutants followed by whole genome sequencing

To select for spontaneous resistant mutants, WT *Mabs* (1 × 10^8^ CFU) was then plated on LB agar supplemented with three different inhibitory concentrations of ETH (128, 150, and 200 μg/ml), which represents 2-, 2.3-, and 3.1-times MIC. Plates were incubated at 37°C for at least 7 days before inspection of the plates. Spontaneous resistant clones were picked and grown in 7H9 broth. ETH resistance was validated in the selected clones using the MIC protocol described above. Genomic DNA was extracted from the clones and submitted to the UC Davis DNA Technologies Core for whole genome sequencing on a NovaSeq platform. Sequence reads were mapped to the *Mabs* ATCC 19977 genome.

### Quantitative PCR (qPCR)

Bacteria at an OD600nm of 0.1 were treated with and without ETH (16 μg/ml). At 3- and 11-hours post-treatment, bacteria were harvested at 3500 rpm for 10 minutes and RNA was extracted using TRIzol (Invitrogen) and purified using the RNeasy Mini Kit (Qiagen) following the manufacturer’s protocol. Total RNA was reversed transcribed to cDNA using the Superscript™ III First-Strand Synthesis System (Invitrogen) following the manufacturer’s protocol. For each primer pair, ten-fold serial dilutions of cDNA were prepared for generation of standard amplification curves on a CFX Connect-Real Time PCR Detection System (Bio-Rad). Fluorescence was detected using the SsoAdvanced Universal SYBR Green Supermix (Bio-Rad). cDNA synthesis reactions without RT were included in parallel to control for genomic DNA contamination. Expression levels were normalized to *sigA* (MAB_3009) before calculating relative expression levels using the delta-delta C_T_ method (2^-ΔΔC^_T_).

### Plasmids

1000bp upstream of *marR*, the *marR* coding sequence, and 200 bp downstream of *marR* were PCR amplified using Q5 DNA Polymerase (New England Biolabs) and cloned into the DraI and HindIII sites of the integrative vector pMV306 (67), generating pRR107. pRR107 was electroporated into Δ*marR* as described above and selected using kanamycin. The *mmpS5-L5* coding sequence along with 200 bp downstream of *mmpL5* were PCR amplified using Q5 DNA Polymerase and cloned into the anhydrotetracycline (ATc) inducible promoter of pUV15LD (56), generating pRR115. pRR115 was electroporated into both WT and Δ*marR* bacteria and selected using Hygromycin B. To check for mycobacterial clones carrying desired plasmids, individual colonies were picked into 10 μl of sterile, nuclease-free water and heat-killed at 80°C for 1 hour. 2 μl of heat-killed bacteria was used as a template for PCR using plasmid-specific primers.

## Funding and Acknowledgements

Sequencing of bacterial mutants and Tn-Seq libraries was carried out by the DNA Technologies and Expression Analysis Core at the UC Davis Genome Center, supported by NIH Shared Instrumentation Grant 1S10OD010786-01. Bioinformatics support was provided by the UC Davis Bioinformatics Core. We would like to thank the Chao Qi lab (Northwestern University) for providing clinical strains, Kayla Dinshaw (Stanley Lab, University of California, Berkeley) for assistance with bacterial genetics, Heran Darwin (New York University) for reading a draft version of this manuscript, and the Stanley and Cox Labs for helpful discussions. Funding was provided by NIH 1R01AI143722 to SAS, NSF GRFP and the UC Berkeley Chancellor’s fellowship to RR.

---

**Figure S1.**
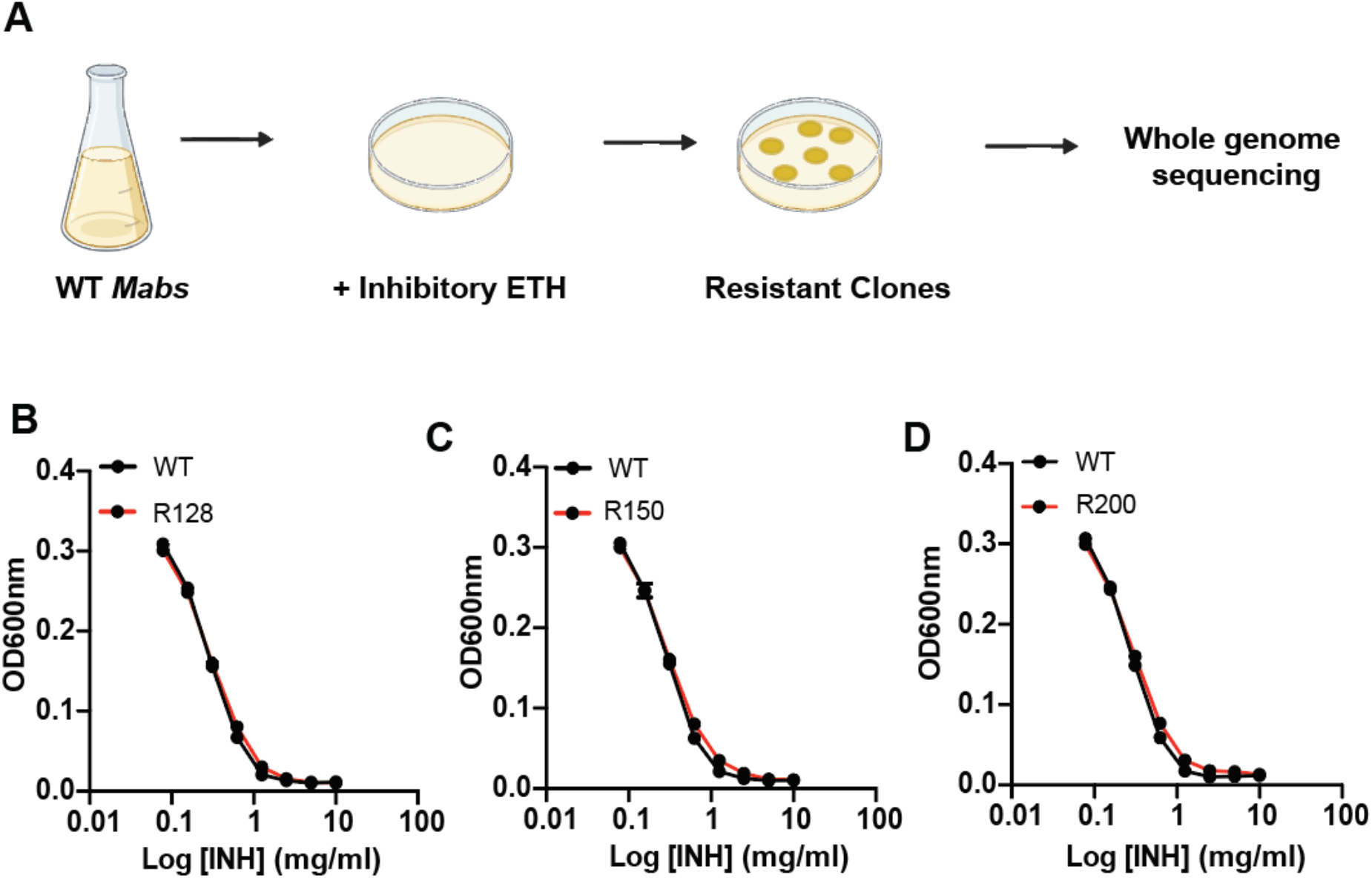
Spontaneous ETH resistance does not confer cross-resistance to INH. (A) WT *Mabs* was plated in the presence of inhibitory ETH to generate resistant bacteria, which were then analyzed by whole genome sequencing. (B-D) Isoniazid (INH) dose-response curves generated against isolated ETH-resistant clones (R128, R150, and R200). Experiments are representative of at least two biological replicates. Error bars represent standard deviation.

**Figure S2.**
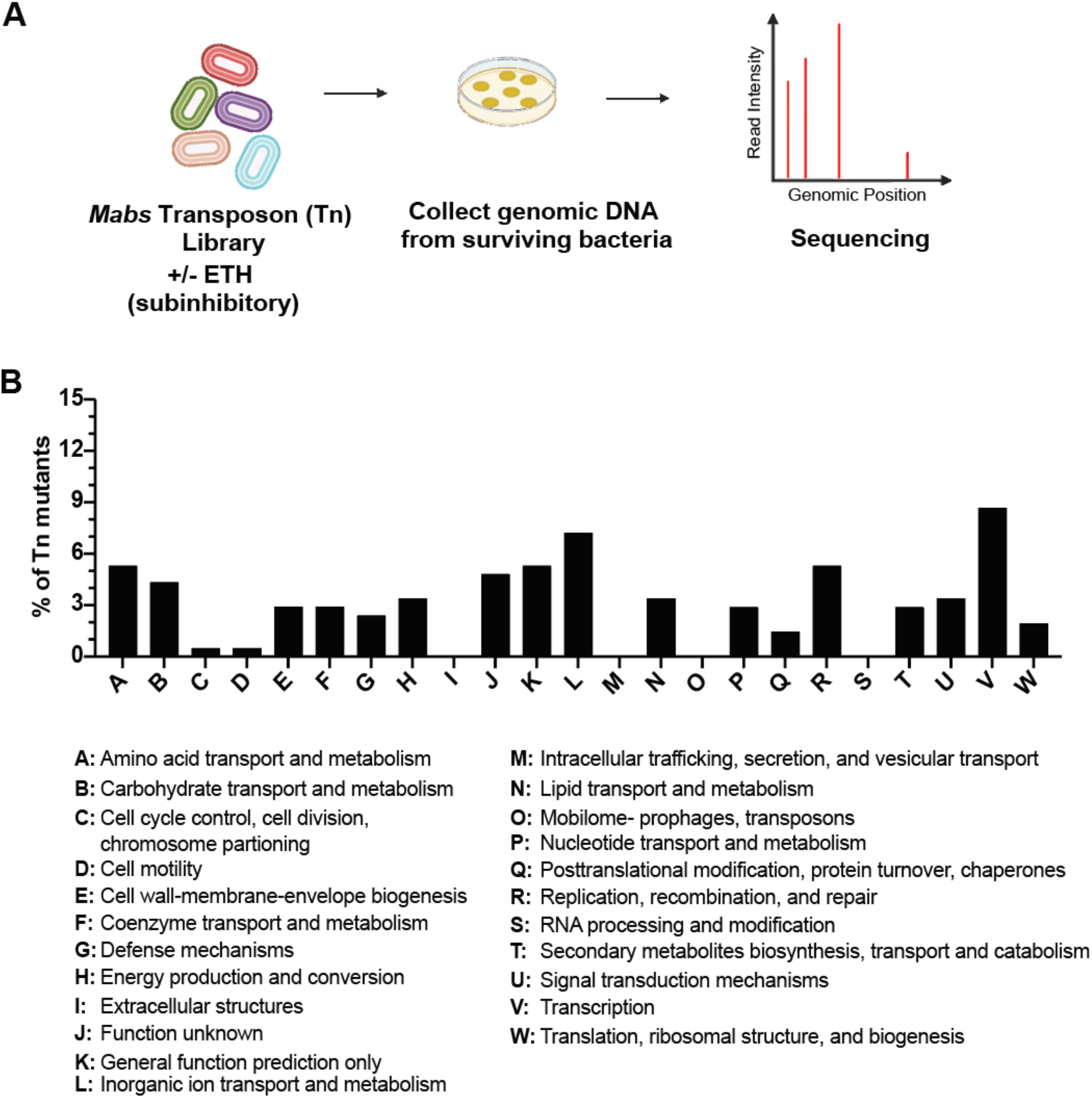
Cluster of Orthologous Group (COG) analysis of transposon mutants exposed to ETH. (A) A library of transposon mutants was treated with and without ETH. Surviving bacteria were then plated, genomic DNA collected, and transposon-enriched sequences analyzed by whole genome sequencing. (B) Percentage of transposon mutants represented in the indicated COG categories. Gene hits without any COG annotations are not shown.

**Figure S3.**
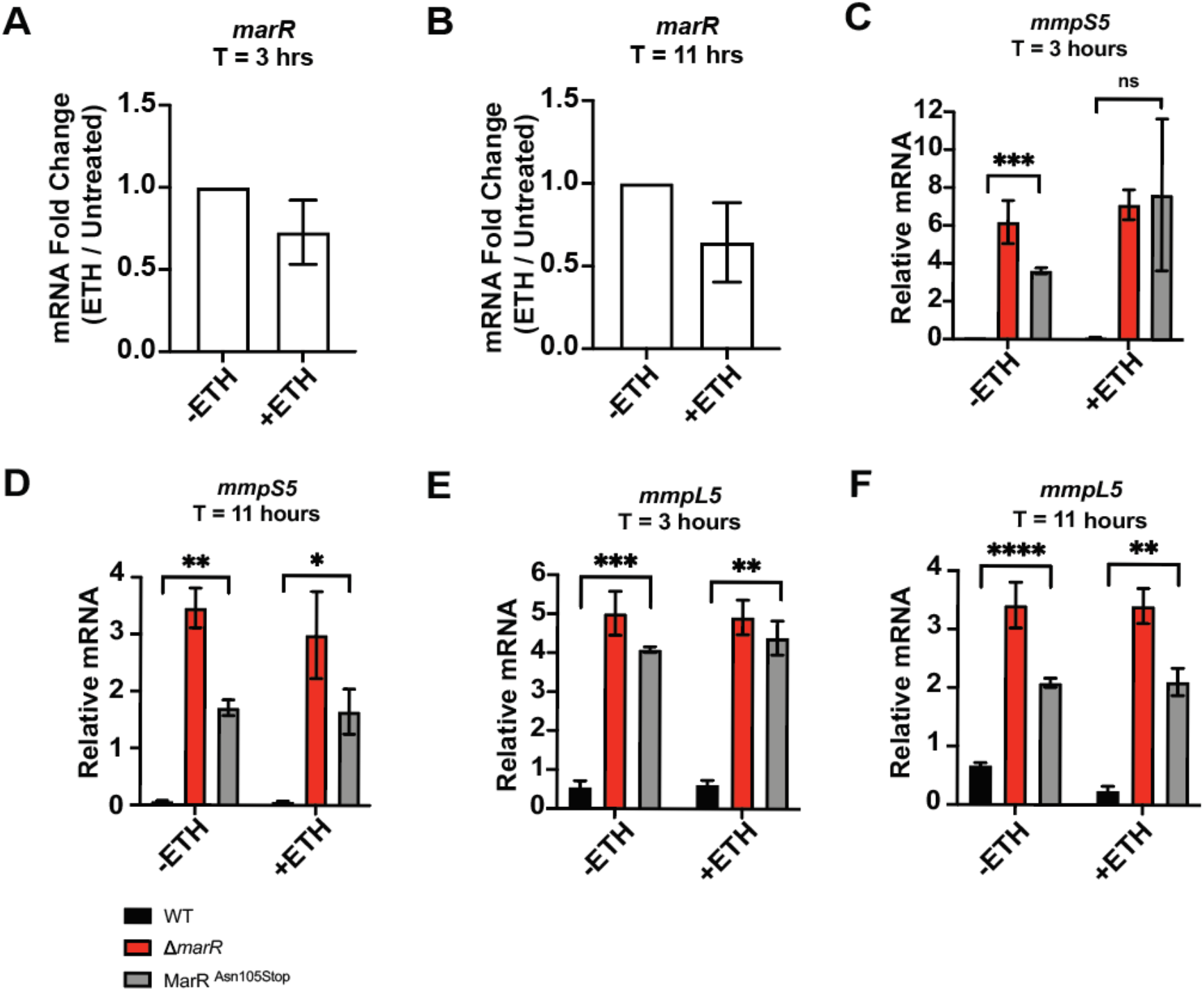
*marR* is not induced in the presence of ETH. (A-B) Gene expression levels of *marR* in WT *Mabs* in the presence and absence of ETH after 3 (A) and 11 hours (B) of exposure. (C-F) Gene expression levels of *mmpS5* and *mmpL5* in WT, Δ*marR*, and MarR^Asn105Stop^ bacteria in the presence and absence of ETH at the indicated time points. Experiments are representative of at least two biological replicates. Error bars represent standard deviation. no significance (ns); *p* < 0.05 (*); *p* < 0.01 (**); *p* < 0.001 (***); *p* < 0.0001 (****).

**Figure S4.**
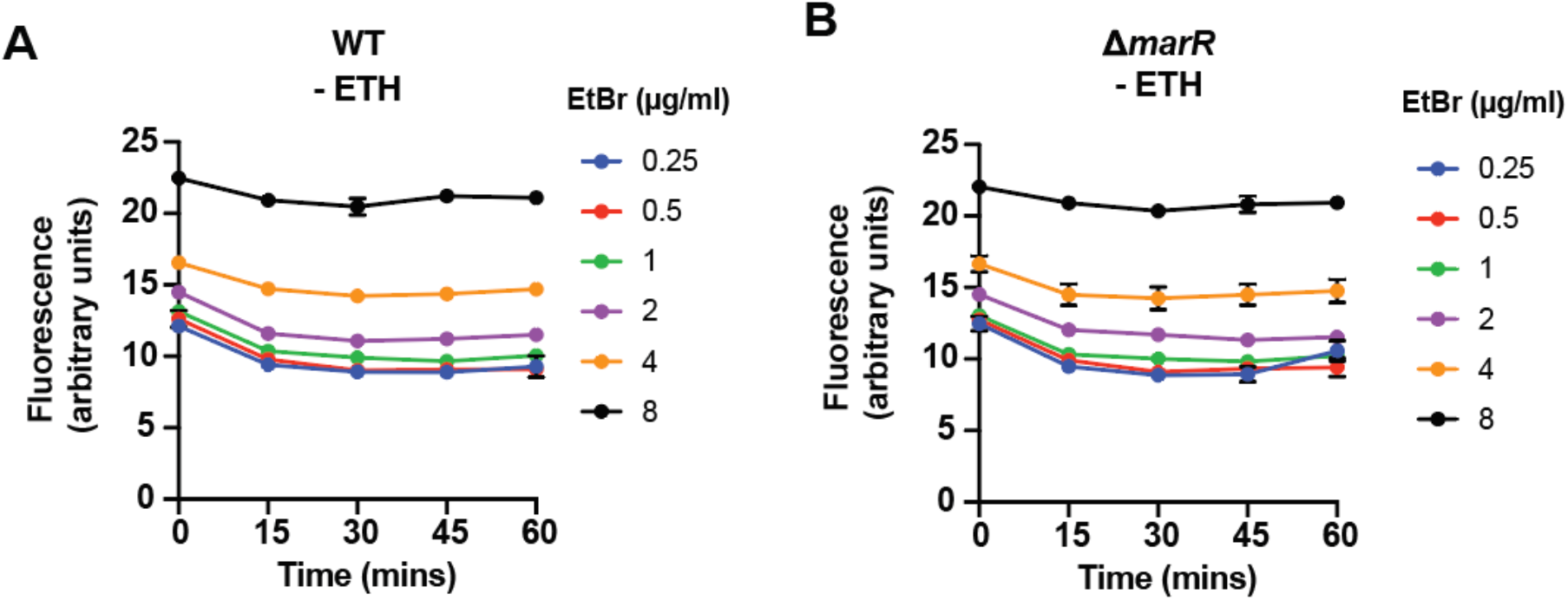
Loss of MarR activity does not lead to changes in Ethidium Bromide (EtBr) accumulation. WT (A) and Δ*marR* (B) *Mabs* were treated with two-fold dilutions of EtBr (0.25 – 8 μg/ml) in PBS supplemented with 0.4% glucose (pH 7.4) and fluorescence measured for 60 minutes at 37°C (Excitation wavelength: 530 nm; Emission wavelength: 585 nm).

**Figure S5.**
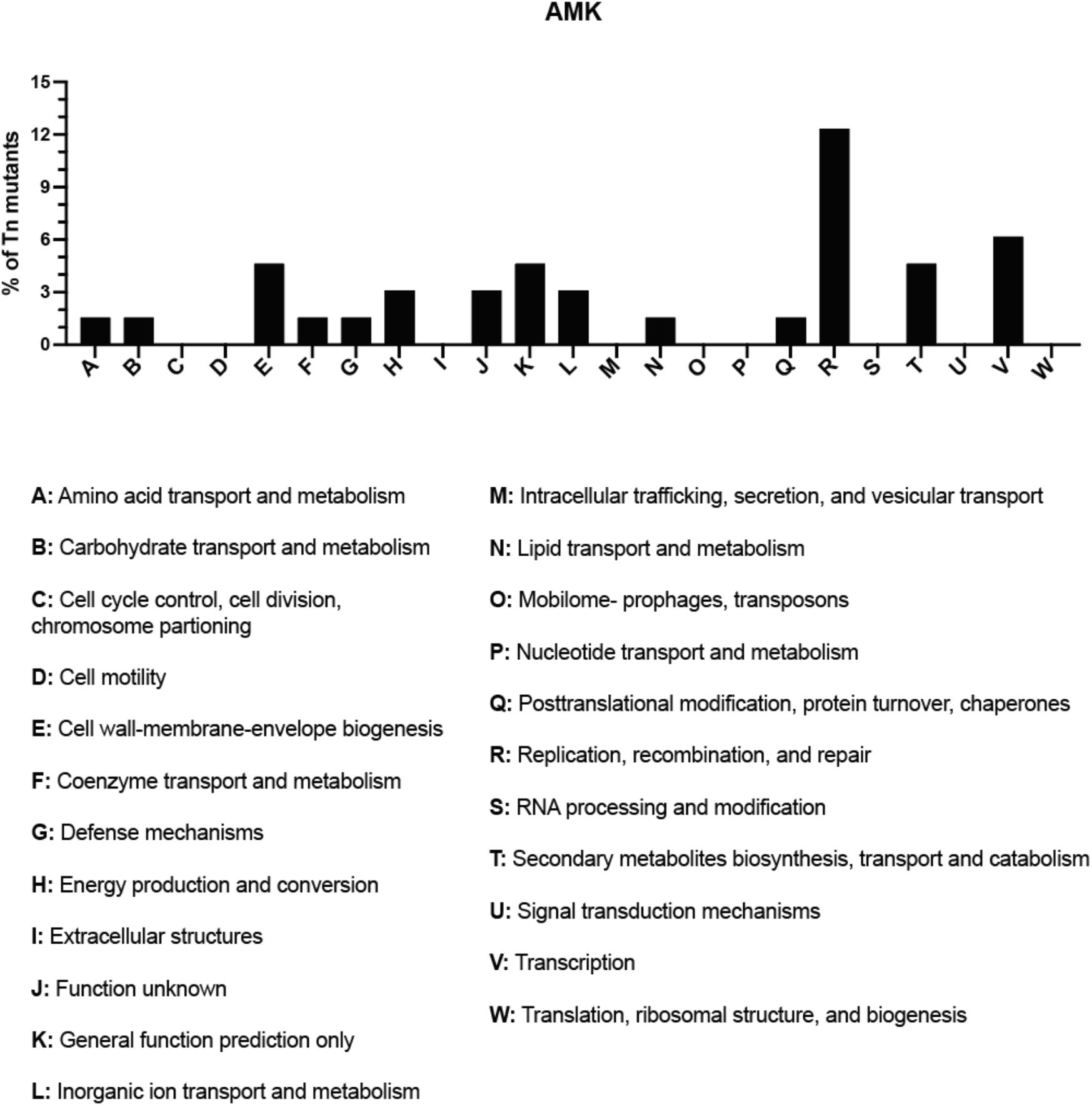
COG analysis of transposon mutants exposed to AMK. Percentage of transposon mutants exposed to AMK represented in the indicated COG categories. Gene hits without any COG annotations are not shown.

**Figure S6.**
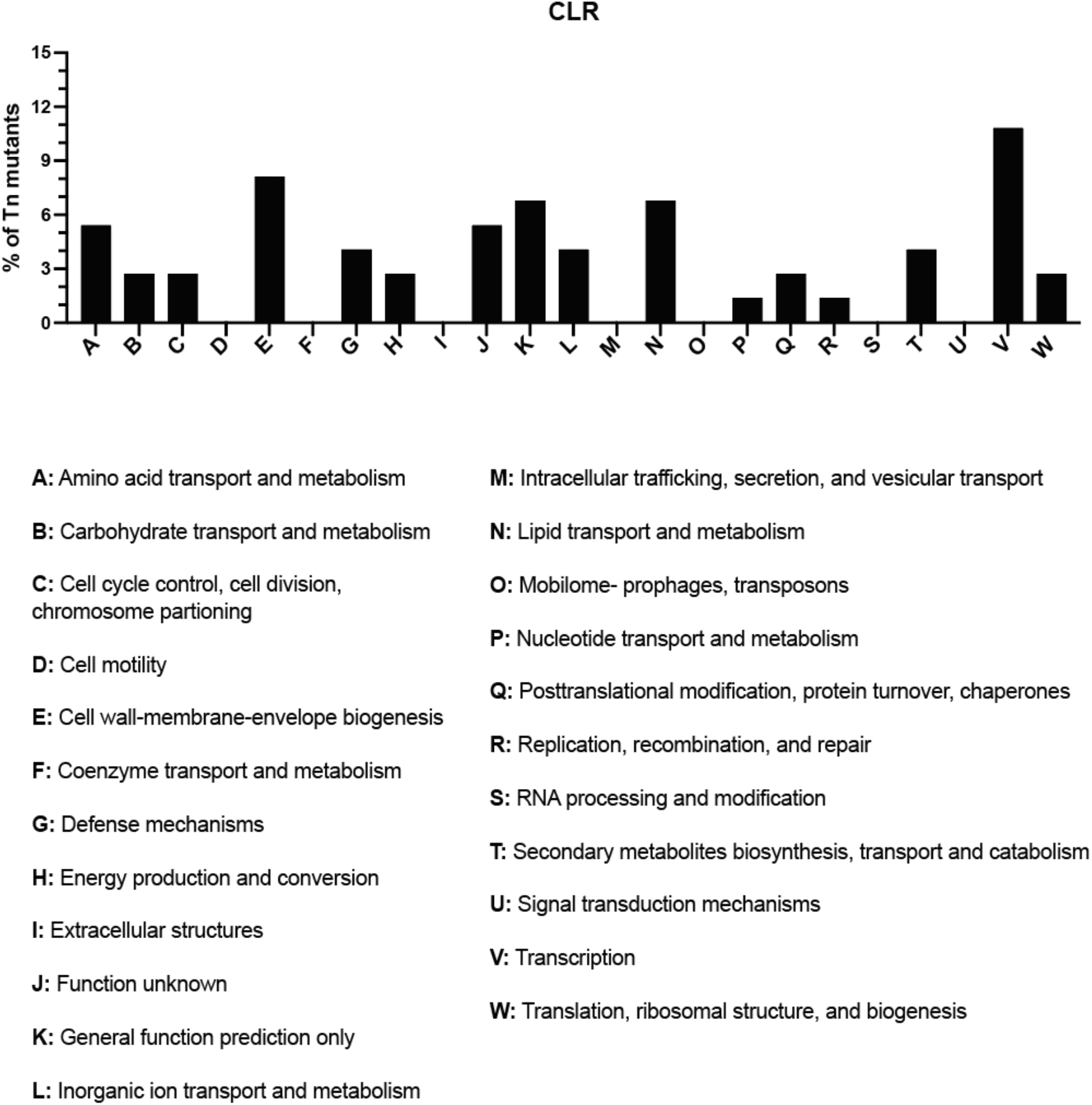
COG analysis of transposon mutants exposed to CLR. Percentage of transposon mutants exposed to CLR represented in the indicated COG categories. Gene hits without any COG annotations are not shown.

**Table S1.**
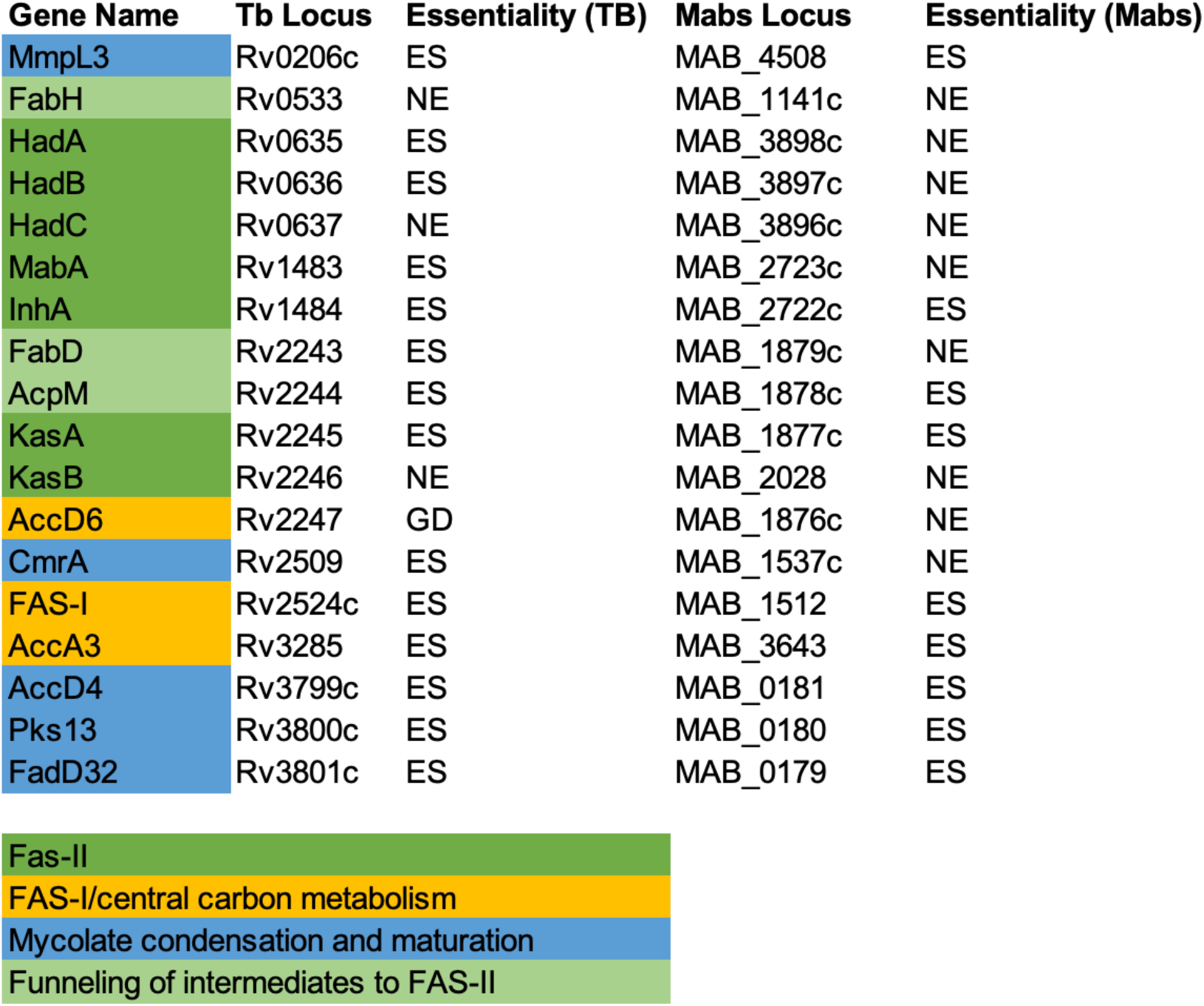
Most mycolic acid biosynthetic genes are essential in both *Mtb* and *Mabs*. ES, essential; NE; not essential; GD, growth defect

| Gene Name | Tb Locus | Essentiality (TB) | Mabs Locus | Essentiality (Mabs) |
| --- | --- | --- | --- | --- |
| MmpL3 | Rv0206c | ES | MAB_4508 | ES |
| FabH | Rv0533 | NE | MAB_1141c | NE |
| HadA | Rv0635 | ES | MAB_3898c | NE |
| HadB | Rv0636 | ES | MAB_3897c | NE |
| HadC | Rv0637 | NE | MAB_3896c | NE |
| MabA | Rv1483 | ES | MAB_2723c | NE |
| InhA | Rv1484 | ES | MAB_2722c | ES |
| FabD | Rv2243 | ES | MAB_1879c | NE |
| AcpM | Rv2244 | ES | MAB_1878c | ES |
| KasA | Rv2245 | ES | MAB_1877c | ES |
| KasB | Rv2246 | NE | MAB_2028 | NE |
| AccD6 | Rv2247 | GD | MAB_1876c | NE |
| CmrA | Rv2509 | ES | MAB_1537c | NE |
| FAS-I | Rv2524c | ES | MAB_1512 | ES |
| AccA3 | Rv3285 | ES | MAB_3643 | ES |
| AccD4 | Rv3799c | ES | MAB_0181 | ES |
| Pks13 | Rv3800c | ES | MAB_0180 | ES |
| FadD32 | Rv3801c | ES | MAB_0179 | ES |

|  |
| --- |
| Fas-II |
| FAS-I/central carbon metabolism |
| Mycolate condensation and maturation |
| Funneling of intermediates to FAS-II |

**Table S3.** Comparison of MIC values in WT and Δ*marR Mabs* to functionally diverse antibiotics.

| Antibiotic | WT MIC (µg/ml) | $\Delta marR$ MIC (µg/ml) |
| --- | --- | --- |
| Amikacin | 2 | 2 |
| Bedaquiline | 0.25 | 0.25 |
| Cefoxitin | 4 | 4 |
| Clofazimine | >16 | >16 |
| Ethambutol | 32 | 32 |
| Isoniazid | 1250 | 1250 |
| Thiacetazone | > 500 | >500 |
| Imipenem | 4 | 4 |
| *p*-aminosalicylic acid | > 160 | > 160 |
| Moxifloxacin | 1 | 1 |
| Doxycycline | > 64 | > 64 |
| Clarithromycin | 4 | 4 |

**Table S4.** ETH MIC values in WT *Mabs* in the presence of each indicated antibiotic. AMK: Amikacin; CLR: Clarithromycin; MFX: Moxifloxacin; CEF: Cefoxitin; IPM: Imipenem.

| Antibiotic | ETH MIC (µg/ml) |
| --- | --- |
| AMK | 8 |
| CLR | 8 |
| MFX | 8 |
| CEF | 4 |
| IPM | 4 |

## References

1. Tortoli E, Kohl TA, Trovato A, Baldan R, Campana S, Cariani L, Colombo C, Costa D, Cristadoro S, Di Serio MC, Manca A, Pizzamiglio G, Rancoita PMV, Rossolini GM, Taccetti G, Teri A, Niemann S, Cirillo DM. 2017. Mycobacterium abscessus in patients with cystic fibrosis: low impact of inter-human transmission in Italy. Eur Respir J 50:1602525.

2. Schuurbiers MMF, Bruno M, Zweijpfenning SMH, Magis-Escurra C, Boeree M, Netea MG, van Ingen J, van de Veerdonk F, Hoefsloot W. 2020. Immune defects in patients with pulmonary Mycobacterium abscessus disease without cystic fibrosis. ERJ Open Res 6:00590–02020.

3. Daniel-Wayman S, Adjemian J, Rebecca Prevots D. 2019. Epidemiology of nontuberculous Mycobacterial pulmonary disease (NTM PD) in the USA, p. 145–161. In Nontuberculous Mycobacterial Disease. Springer International Publishing, Cham.

4. Kendall BA, Winthrop KL. 2013. Update on the epidemiology of pulmonary nontuberculous mycobacterial infections. Semin Respir Crit Care Med 34:87–94.

5. Lai C-C, Tan C-K, Chou C-H, Hsu H-L, Liao C-H, Huang Y-T, Yang P-C, Luh K-T, Hsueh P-R. 2010. Increasing incidence of nontuberculous mycobacteria, Taiwan, 2000–2008. Emerg Infect Dis 16:1047b–11048.

6. Gardner AI, McClenaghan E, Saint G, McNamara PS, Brodlie M, Thomas MF. 2019. Epidemiology of nontuberculous mycobacteria infection in children and young people with cystic fibrosis: Analysis of UK cystic fibrosis Registry. Clin Infect Dis 68:731– 737.

7. Nie W, Duan H, Huang H, Lu Y, Bi D, Chu N. 2014. Species identification of Mycobacterium abscessus subsp. abscessus and Mycobacterium abscessus subsp. bolletii using rpoB and hsp65, and susceptibility testing to eight antibiotics. Int J Infect Dis 25:170–174.

8. Dartois V, Dick T. 2022. Drug development challenges in nontuberculous mycobacterial lung disease: TB to the rescue. J Exp Med 219.

9. Gumbo T, Cirrincione K, Srivastava S. 2020. Repurposing drugs for treatment of Mycobacterium abscessus: a view to a kill. J Antimicrob Chemother 75:1212–1217.

10. Lee M-R, Sheng W-H, Hung C-C, Yu C-J, Lee L-N, Hsueh P-R. 2015. Mycobacterium abscessus Complex Infections in Humans. Emerg Infect Dis 21:1638–1646.

11. Chandrashekaran S, Escalante P, Kennedy CC. 2017. Mycobacterium abscessus disease in lung transplant recipients: Diagnosis and management. J Clin Tuberc Other Mycobact Dis 9:10–18.

12. Floto RA, Olivier KN, Saiman L, Daley CL, Herrmann J-L, Nick JA, Noone PG, Bilton D, Corris P, Gibson RL, Hempstead SE, Koetz K, Sabadosa KA, Sermet-Gaudelus I, Smyth AR, van Ingen J, Wallace RJ, Winthrop KL, Marshall BC, Haworth CS, US Cystic Fibrosis Foundation and European Cystic Fibrosis Society. 2016. US Cystic Fibrosis Foundation and European Cystic Fibrosis Society consensus recommendations for the management of non-tuberculous mycobacteria in individuals with cystic fibrosis. Thorax 71 Suppl 1:i1–22.

13. Griffith DE. 2019. Mycobacterium abscessus and antibiotic resistance: Same as it ever was. Clin Infect Dis. Oxford University Press (OUP).

14. Rudra P, Hurst-Hess K, Lappierre P, Ghosh P. 2018. High levels of intrinsic tetracycline resistance in Mycobacterium abscessus are conferred by a tetracycline-modifying monooxygenase. Antimicrob Agents Chemother 62.

15. Soroka D, Dubée V, Soulier-Escrihuela O, Cuinet G, Hugonnet J-E, Gutmann L, Mainardi J-L, Arthur M. 2014. Characterization of broad-spectrum Mycobacterium abscessus class A β-lactamase. J Antimicrob Chemother 69:691–696.

16. Hurst-Hess K, Rudra P, Ghosh P. 2017. Mycobacterium abscessus WhiB7 regulates a species-specific repertoire of genes to confer extreme antibiotic resistance. Antimicrob Agents Chemother 61.

17. Luthra S, Rominski A, Sander P. 2018. The role of antibiotic-target-modifying and antibiotic-modifying enzymes in Mycobacterium abscessus drug resistance. Front Microbiol 9:2179.

18. Viljoen A, Dubois V, Girard-Misguich F, Blaise M, Herrmann J-L, Kremer L. 2017. The diverse family of MmpL transporters in mycobacteria: from regulation to antimicrobial developments. Mol Microbiol 104:889–904.

19. Marrakchi H, Lanéelle M-A, Daffé M. 2014. Mycolic acids: structures, biosynthesis, and beyond. Chem Biol 21:67–85.

20. Armitige LY, Jagannath C, Wanger AR, Norris SJ. 2000. Disruption of the genes encoding antigen 85A and antigen 85B of Mycobacterium tuberculosis H37Rv: effect on growth in culture and in macrophages. Infect Immun 68:767–778.

21. Liu J, Barry CE III, Besra GS, Nikaido H. 1996. Mycolic acid structure determines the fluidity of the Mycobacterial cell wall. J Biol Chem 271:29545–29551.

22. Rozwarski DA, Grant GA, Barton DH, Jacobs WR Jr, Sacchettini JC. 1998. Modification of the NADH of the isoniazid target (InhA) from Mycobacterium tuberculosis. Science 279:98–102.

23. Vilchèze C, Jacobs WR Jr. 2014. Resistance to Isoniazid and Ethionamide in Mycobacterium tuberculosis: Genes, Mutations, and Causalities. Microbiol Spectr 2:MGM2-0014–2013.

24. Li G, Lian L-L, Wan L, Zhang J, Zhao X, Jiang Y, Zhao L-L, Liu H, Wan K. 2013. Antimicrobial susceptibility of standard strains of nontuberculous mycobacteria by microplate Alamar Blue assay. PLoS One 8:e84065.

25. Reingewertz TH, Meyer T, McIntosh F, Sullivan J, Meir M, Chang Y-F, Behr MA, Barkan D. 2020. Differential sensitivity of mycobacteria to isoniazid is related to differences in KatG-mediated enzymatic activation of the drug. Antimicrob Agents Chemother 64.

26. Grzegorzewicz AE, Eynard N, Quémard A, North EJ, Margolis A, Lindenberger JJ, Jones V, Korduláková J, Brennan PJ, Lee RE, Ronning DR, McNeil MR, Jackson M. 2015. Covalent modification of the Mycobacterium tuberculosis FAS-II dehydratase by Isoxyl and Thiacetazone. ACS Infect Dis 1:91–97.

27. Halloum I, Viljoen A, Khanna V, Craig D, Bouchier C, Brosch R, Coxon G, Kremer L. 2017. Resistance to thiacetazone derivatives active against Mycobacterium abscessus involves mutations in the MmpL5 transcriptional repressor MAB_4384. Antimicrob Agents Chemother 61.

28. Costa-Gouveia J, Pancani E, Jouny S, Machelart A, Delorme V, Salzano G, Iantomasi R, Piveteau C, Queval CJ, Song O-R, Flipo M, Deprez B, Saint-André J-P, Hureaux J, Majlessi L, Willand N, Baulard A, Brodin P, Gref R. 2017. Combination therapy for tuberculosis treatment: pulmonary administration of ethionamide and booster co-loaded nanoparticles. Sci Rep 7.

29. Baulard AR, Betts JC, Engohang-Ndong J, Quan S, McAdam RA, Brennan PJ, Locht C, Besra GS. 2000. Activation of the pro-drug ethionamide is regulated in mycobacteria. J Biol Chem 275:28326–28331.

30. Wang F, Langley R, Gulten G, Dover LG, Besra GS, Jacobs WR Jr, Sacchettini JC. 2007. Mechanism of thioamide drug action against tuberculosis and leprosy. J Exp Med 204:73–78.

31. Ushtanit A, Kulagina E, Mikhailova Y, Makarova M, Safonova S, Zimenkov D. 2022. Molecular determinants of ethionamide resistance in clinical isolates of Mycobacterium tuberculosis. Antibiotics (Basel) 11:133.

32. Cain AK, Barquist L, Goodman AL, Paulsen IT, Parkhill J, van Opijnen T. 2020. A decade of advances in transposon-insertion sequencing. Nat Rev Genet 21:526– 540.

33. Gallagher LA, Shendure J, Manoil C. 2011. Genome-scale identification of resistance functions in Pseudomonas aeruginosa using Tn-seq. MBio 2:e00315–10.

34. Rajagopal M, Martin MJ, Santiago M, Lee W, Kos VN, Meredith T, Gilmore MS, Walker S. 2016. Multidrug intrinsic resistance factors in Staphylococcus aureus identified by profiling fitness within high-diversity transposon libraries. MBio 7.

35. Philalay JS, Palermo CO, Hauge KA, Rustad TR, Cangelosi GA. 2004. Genes Required for Intrinsic Multidrug Resistance in Mycobacterium avium. Antimicrob Agents Chemother 48:3412–3418.

36. Akusobi C, Benghomari BS, Zhu J, Wolf ID, Singhvi S, Dulberger CL, Ioerger TR, Rubin EJ. 2022. Transposon mutagenesis in Mycobacterium abscessus identifies an essential penicillin-binding protein involved in septal peptidoglycan synthesis and antibiotic sensitivity. Elife 11.

37. Rifat D, Chen L, Kreiswirth BN, Nuermberger EL. 2021. Genome-wide essentiality analysis of Mycobacterium abscessus by saturated transposon Mutagenesis and deep sequencing. MBio 12:e0104921.

38. Sullivan MR, McGowen K, Liu Q, Akusobi C, Young DC, Mayfield JA, Raman S, Wolf ID, Branch Moody D, Aldrich CC, Muir A, Rubin EJ. 2022. Cell envelope remodeling requires high concentrations of biotin during Mycobacterium abscessus model lung infection. bioRxiv.

39. Richard M, Gutiérrez AV, Viljoen A, Rodriguez-Rincon D, Roquet-Baneres F, Blaise M, Everall I, Parkhill J, Floto RA, Kremer L. 2019. Mutations in the MAB_2299c TetR regulator confer cross-resistance to clofazimine and bedaquiline in Mycobacterium abscessus. Antimicrob Agents Chemother 63.

40. Sassetti CM, Boyd DH, Rubin EJ. 2001. Comprehensive identification of conditionally essential genes in mycobacteria. Proc Natl Acad Sci U S A 98:12712–12717.

41. Ripoll F, Pasek S, Schenowitz C, Dossat C, Barbe V, Rottman M, Macheras E, Heym B, Herrmann J-L, Daffé M, Brosch R, Risler J-L, Gaillard J-L. 2009. Non mycobacterial virulence genes in the genome of the emerging pathogen Mycobacterium abscessus. PLoS One 4:e5660.

42. Medjahed H, Singh AK. 2010. Genetic manipulation of Mycobacterium abscessus. Curr Protoc Microbiol Chapter 10:Unit 10D.2.

43. Long JE, DeJesus M, Ward D, Baker RE, Ioerger T, Sassetti CM. 2015. Identifying essential genes in Mycobacterium tuberculosis by global phenotypic profiling. Methods Mol Biol 1279:79–95.

44. DeJesus MA, Ambadipudi C, Baker R, Sassetti C, Ioerger TR. 2015. TRANSIT--A software tool for Himar1 TnSeq analysis. PLoS Comput Biol 11:e1004401.

45. Minato Y, Gohl DM, Thiede JM, Chacón JM, Harcombe WR, Maruyama F, Baughn AD. 2019. Genomewide assessment of Mycobacterium tuberculosis conditionally essential metabolic pathways. mSystems 4.

46. Daffé M, Quémard A, Marrakchi H. 2017. Mycolic Acids: From Chemistry to Biology, p. 1–36. In Geiger, O (ed.), Biogenesis of Fatty Acids, Lipids and Membranes. Springer International Publishing, Cham.

47. Nolan CM. 2003. Isoniazid for latent tuberculosis infection: approaching 40 and reaching its prime. Am J Respir Crit Care Med. American Thoracic Society.

48. Vilchèze C, Wang F, Arai M, Hazbón MH, Colangeli R, Kremer L, Weisbrod TR, Alland D, Sacchettini JC, Jacobs WR Jr. 2006. Transfer of a point mutation in Mycobacterium tuberculosis inhA resolves the target of isoniazid. Nat Med 12:1027– 1029.

49. Heysell SK, Pholwat S, Mpagama SG, Pazia SJ, Kumburu H, Ndusilo N, Gratz J, Houpt ER, Kibiki GS. 2015. Sensititre MycoTB plate compared to Bactec MGIT 960 for first- and second-line antituberculosis drug susceptibility testing in Tanzania: a call to operationalize MICs. Antimicrob Agents Chemother 59:7104–7108.

50. Auclair B, Nix DE, Adam RD, James GT, Peloquin CA. 2001. Pharmacokinetics of ethionamide administered under fasting conditions or with orange juice, food, or antacids. Antimicrob Agents Chemother 45:810–814.

51. Cuthbertson L, Nodwell JR. 2013. The TetR family of regulators. Microbiol Mol Biol Rev 77:440–475.

52. Morlock GP, Metchock B, Sikes D, Crawford JT, Cooksey RC. 2003. ethA, inhA, and katG loci of ethionamide-resistant clinical Mycobacterium tuberculosis isolates. Antimicrob Agents Chemother 47:3799–3805.

53. Cohen SP, McMurry LM, Hooper DC, Wolfson JS, Levy SB. 1989. Cross-resistance to fluoroquinolones in multiple-antibiotic-resistant (Mar) Escherichia coli selected by tetracycline or chloramphenicol: decreased drug accumulation associated with membrane changes in addition to OmpF reduction. Antimicrob Agents Chemother 33:1318–1325.

54. Kotecka K, Kawalek A, Kobylecki K, Bartosik AA. 2021. The MarR-type regulator PA3458 is involved in osmoadaptation control in Pseudomonas aeruginosa. Int J Mol Sci 22:3982.

55. Ziha-Zarifi I, Llanes C, Köhler T, Pechere J-C, Plesiat P. 1999. In vivo emergence of multidrug-resistant mutants of Pseudomonas aeruginosa overexpressing the active efflux system MexA-MexB-OprM. Antimicrob Agents Chemother 43:287–291.

56. Ehrt S, Guo XV, Hickey CM, Ryou M, Monteleone M, Riley LW, Schnappinger D. 2005. Controlling gene expression in mycobacteria with anhydrotetracycline and Tet repressor. Nucleic Acids Res 33:e21.

57. Blair JMA, Richmond GE, Piddock LJV. 2014. Multidrug efflux pumps in Gram-negative bacteria and their role in antibiotic resistance. Future Microbiol 9:1165– 1177.

58. Rodrigues L, Ramos J, Couto I, Amaral L, Viveiros M. 2011. Ethidium bromide transport across Mycobacterium smegmatis cell-wall: correlation with antibiotic resistance. BMC Microbiol 11:35.

59. Kapopoulou A, Lew JM, Cole ST. 2011. The MycoBrowser portal: a comprehensive and manually annotated resource for mycobacterial genomes. Tuberculosis (Edinb) 91:8–13.

60. Chubiz LM, Rao CV. 2010. Aromatic acid metabolites of Escherichia coli K-12 can induce the marRAB operon. J Bacteriol 192:4786–4789.

61. Sharma P, Haycocks JRJ, Middlemiss AD, Kettles RA, Sellars LE, Ricci V, Piddock LJV, Grainger DC. 2017. The multiple antibiotic resistance operon of enteric bacteria controls DNA repair and outer membrane integrity. Nat Commun 8:1444.

62. Altschul SF, Gish W, Miller W, Myers EW, Lipman DJ. 1990. Basic local alignment search tool. J Mol Biol 215:403–410.

63. Melly G, Purdy GE. 2019. MmpL Proteins in Physiology and Pathogenesis of M. tuberculosis. Microorganisms 7:70.

64. Wells RM, Jones CM, Xi Z, Speer A, Danilchanka O, Doornbos KS, Sun P, Wu F, Tian C, Niederweis M. 2013. Discovery of a siderophore export system essential for virulence of Mycobacterium tuberculosis. PLoS Pathog 9:e1003120.

65. Mukherjee D, Wu M-L, Teo JWP, Dick T. 2017. Vancomycin and clarithromycin show synergy against Mycobacterium abscessus in vitro. Antimicrob Agents Chemother 61.

66. Murphy KC, Nelson SJ, Nambi S, Papavinasasundaram K, Baer CE, Sassetti CM. 2018. ORBIT: A new paradigm for genetic engineering of Mycobacterial chromosomes. MBio 9.

67. Andreu N, Zelmer A, Fletcher T, Elkington PT, Ward TH, Ripoll J, Parish T, Bancroft GJ, Schaible U, Robertson BD, Wiles S. 2010. Optimisation of bioluminescent reporters for use with mycobacteria. PLoS One 5:e10777.

